## Supplementary information for "A conserved membrane protein negatively regulates Mce1 complexes in mycobacteria"

**Affiliations:**

**This PDF file includes:**

Supplementary Figures 1 to 8
Supplementary Tables 1 to 6
Supplementary References 1 to 13

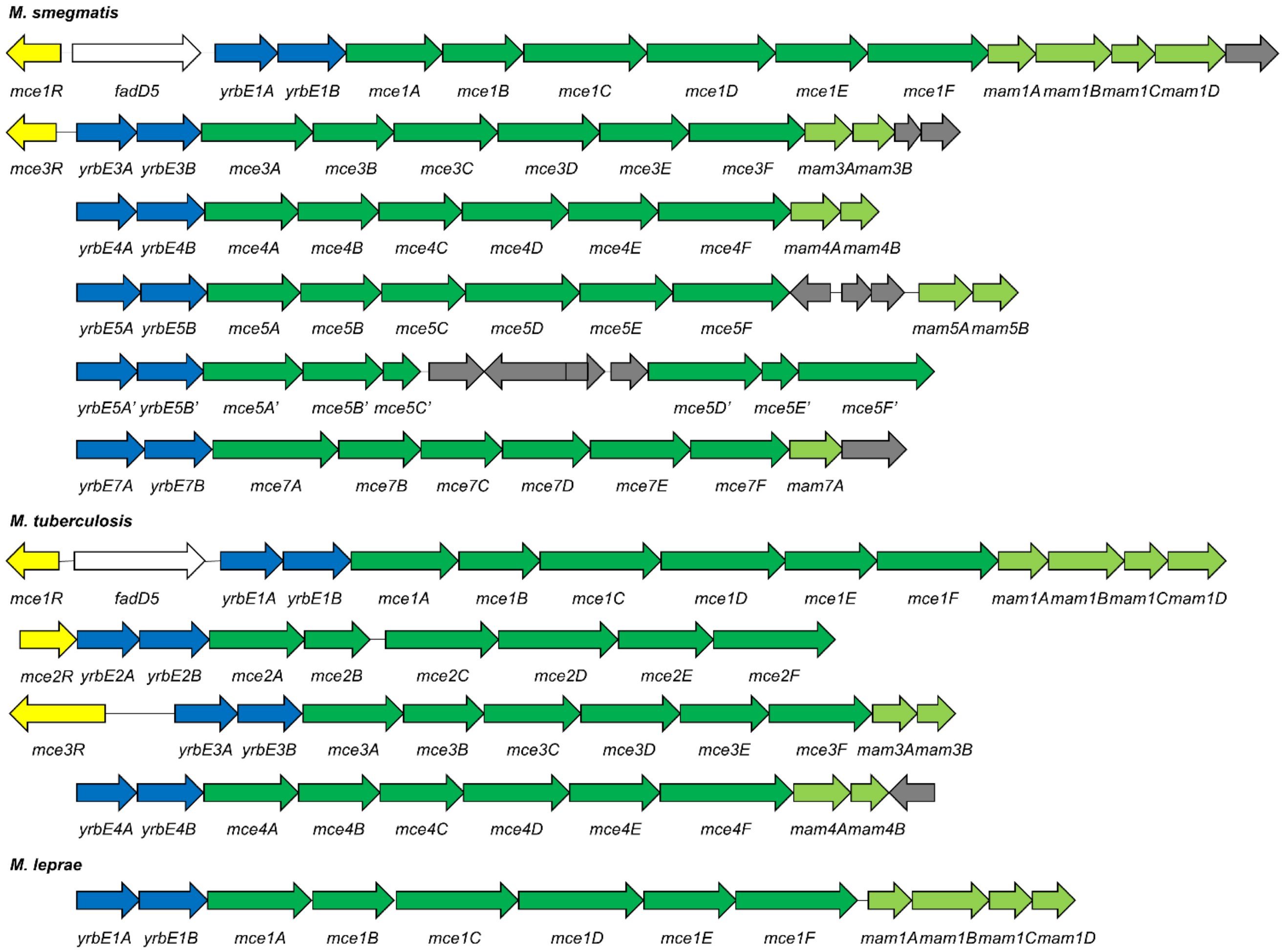


**Supplementary Figure 1. A schematic representation of the *mce* genetic loci in *M. smegmatis*, *M. tuberculosis* and *M. leprae*.** Each straight arrow represents an individual gene. Lengths of the arrows are scaled to the lengths of the genes. The adjacent gene that encodes the transcriptional regulator of some operons is colored in *yellow*. *yrbE* genes are colored in *blue*, *mce* genes *dark green*, *mam* genes *light green*, and unrelated genes *grey*. *mce1* operons in *M. smegmatis* and *M. tuberculosis* contains an additional gene encoding a putative fatty acyl-CoA synthase (*fadD5*, *white*). The *mce5* operon contains an insertion of unrelated genes between *mce5* and *mam5* genes. The *mce5bis* operon (*mce5'*) contains an insertion between *mce5C'* and *mce5D'*.

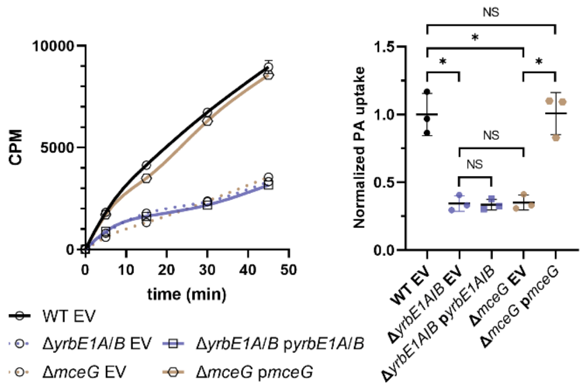


**Supplementary Figure 2. The Δ*mceG* strain can be complemented by exogenous expression of *mceG*.** [$^{14}$C]-palmitic acid uptake profiles and rates of indicated *M. smegmatis* strains. The uptake profile (*left*) shows accumulated radioactivity counts in cells over time, and is representative of at least three independent experiments. Each data point (mean ± standard deviation) represents results from three *technical* replicates. CPM, count per minute. Uptake rates (*right*) are quantified based on [$^{14}$C]-palmitic acid levels after 30 min incubation. The uptake of individual strains is normalized to that of WT cells harboring the empty vector (WT EV). Mean ± standard deviation of three *biological* replicates is shown for each group. Welch's t-test: NS, not significant; * p<0.05; ** p<0.01; *** p<0.001. EV, empty vector (pJEB402). The Δ*yrbE1A*/*B* strain could not be complemented by expressing *yrbE1A*/*B*, very likely due to polar effects on downstream genes. Nevertheless, both the Δ*yrbE1A*/*B* and Δ*mceG* strains represent cells that have lost Mce1 function completely, therefore useful as negative controls.

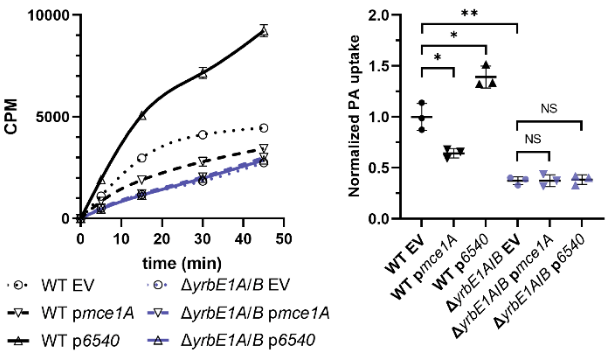


**Supplementary Figure 3. Mce1A and MSMEG_6540 may compete to form distinct Mce1 complexes.** [$^{14}$C]-palmitic acid uptake profiles and rates of indicated *M. smegmatis* strains. The uptake profile (*left*) shows accumulated radioactivity counts in cells over time, and is representative of at least three independent experiments. Each data point (mean ± standard deviation) represents results from three *technical* replicates. CPM, count per minute. Uptake rates (*right*) are quantified based on [$^{14}$C]-palmitic acid levels after 30 min incubation. The uptake of individual strains is normalized to that of WT cells harboring the empty vector (WT EV). Mean ± standard deviation of three *biological* replicates is shown for each group. Welch's t-test: NS, not significant; * p<0.05; ** p<0.01; *** p<0.001. EV, empty vector (pJEB402); *6540*, *MSMEG_6540*.

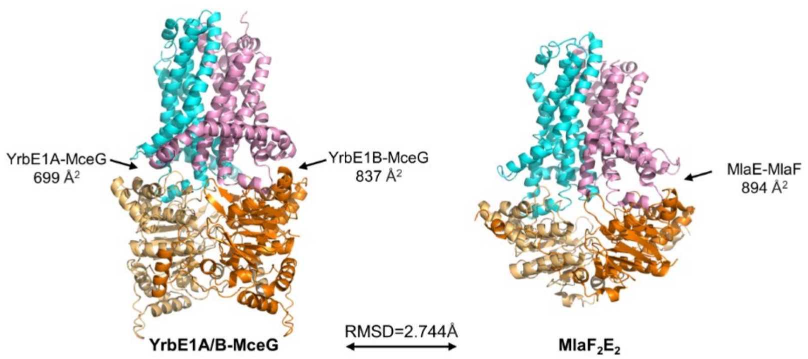


**Supplementary Figure 4. Comparison of the AlphaFold2 structural model of YrbE1A/B-MceG and cryo-electron microscopy structure of MlaFE** (PDB: 7CH6, with MlaB removed)[1]. Left panel, *cyan*: YrbE1A, *pink*: YrbE1B, *light orange* and *orange*: MceG; right panel, *cyan* and *pink*: MlaE, *light orange* and *orange*: MlaF. Calculated interface areas are indicated.

a

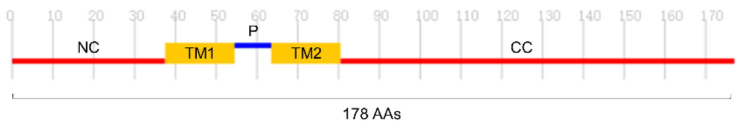


b

```
                1                                                    50
M12418c        ~~~~~~~~~~ ~~~~~~~~~M TAKTEANRKP LFYESGANWI GTSGS...GG
Msmeg_0959     MTEPAGHEPP EAAGMSGVPD ASGAAPAGEV LFYEQGASWA WLLSGPIAGV
Rv0513         ~~~~~~~~~~ ~~~~~~~~~M TPTGDTKPKL LFYEPGASWY WVLTGPLAAV

                51                                                  100
M12418c        RDFHWHFGSG APMQLVVLLI FLVLVSWFVG LQVKPAQIQS SVEPTYGALC
Msmeg_0959     GMAILQRTGG YGYDLWIPLI FLVLVSGFIA IQIKAARIHT SVELTRETLR
Rv0513         SVLLLEISSG AGVGLITPAI FLVMVSAFVA LQVKAARIHT SVELTHDALR

                101                                                 150
M12418c        RGVETILVAE IVKILSGNRE ..FTGVQ~~~ ~~~~~~~~~~ ~~~~~~~~~~
                                                     ▼  ▼      ▼
Msmeg_0959     QGTEIISIDE IVDIYPEASG .....SEVPK WQSARALGEL TGVPRGRTGI
Rv0513         QGTETIRLAE IVKIYPEADG RETSGEEPAK WQSARTLGEL VGVPRGRVGI

                151                                                 200
               ~~~~~~~~~~ ~~~~~~~~~~ ~~~~~~~~~~ ~~~~~~~~~~ ~~~~~~~~~~
M12418c
Msmeg_0959     GLKLTGKRVA QAWARRHRTL RAALQQLVEE RTP~~~~~~~ ~~~~~~~~~~
Rv0513         GLKLTGGRTA QAWARRHQQL RAALTPLVQE RLGPVDSDVA DVNGDDAGPA

                201
               ~
M12418c
               ~
Msmeg_0959
Rv0513         R
```

**Supplementary Figure 5. MSMEG_0959 is a transmembrane protein conserved in mycobacteria. a** A predicted topology model of MSMEG_0959. The length of MSMEG_0959 is 178 amino acids (AAs). Cytoplasmic domains (N-terminal and C-terminal, NC and CC) are colored in *red*, the periplasmic (P) loop in *blue*, and the transmembrane (TM) domains in *yellow*. **b** Sequence alignment of MSMEG_0959 and its homologs in *M. tuberculosis* (Rv0513) and *M. leprae* (ML2418). The protein encoded by *ML2418* lacks the sequence corresponding to a large portion of the C-terminal cytoplasmic domain in MSMEG_0959 due to an early stop codon (only 103 amino acids compared to 178 for MSMEG_0959). The *red* arrowheads indicate residues that were mutated to prolines on the MSMEG_0959 variant (MSMEG_0959$_{AGG3P}$) that no longer competes with MceG for binding to YrbE1B.

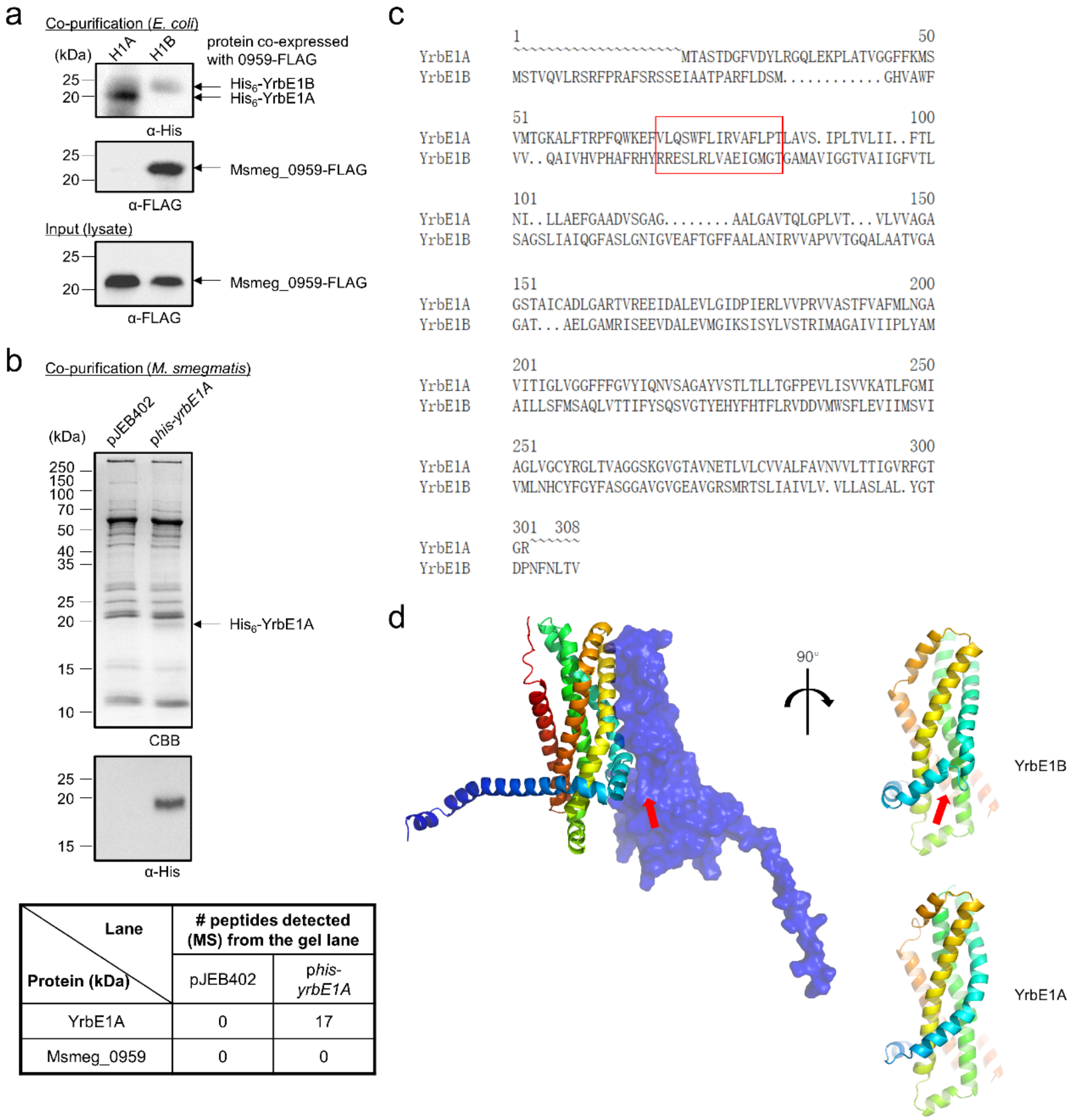


| Lane / Protein (kDa) | # peptides detected (MS) from the gel lane | |
|---|---|---|
| | pJEB402 | *phis-yrbE1A* |
| YrbE1A | 0 | 17 |
| Msmeg_0959 | 0 | 0 |

**Supplementary Figure 6. MSMEG_0959 co-purifies with YrbE1B but not YrbE1A. a** SDS-PAGE and α-His/FLAG immunoblot analyses of proteins affinity-purified from *E. coli* cells expressing His-tagged YrbE1A (H1A) or YrbE1B (H1B) together with FLAG-tagged MSMEG_0959. **b** SDS-PAGE and α-His immunoblot analyses of proteins affinity-purified from *M. smegmatis* cells expressing His-tagged YrbE1A (p*his-yrbE1A)*. pJEB402 was used as the empty vector control. The two entire gel lanes were subjected to MS/MS protein identification. The table shows total numbers of peptides detected for YrbE1A and MSMEG_0959. CBB, Coomassie brilliant blue. **c** Sequence alignment of YrbE1A and YrbE1B. The region of YrbE1B for the putative helix that is required for the interaction of YrbE1B with MSMEG_0959 and the corresponding region of YrbE1A are boxed in *red*. **d** Left panel: An AlphaFold2 model of YrbE1B (*rainbow*) -MSMEG_0959 (shown as *blue* surface). Right panel: comparison of the putative regions facing MSMEG_0959 in YrbE1B and YrbE1A. The red arrows point towards the short helix and its adjoining loop on YrbE1B that likely determines its interaction with MSMEG_0959, which is absent at the same region of YrbE1A.

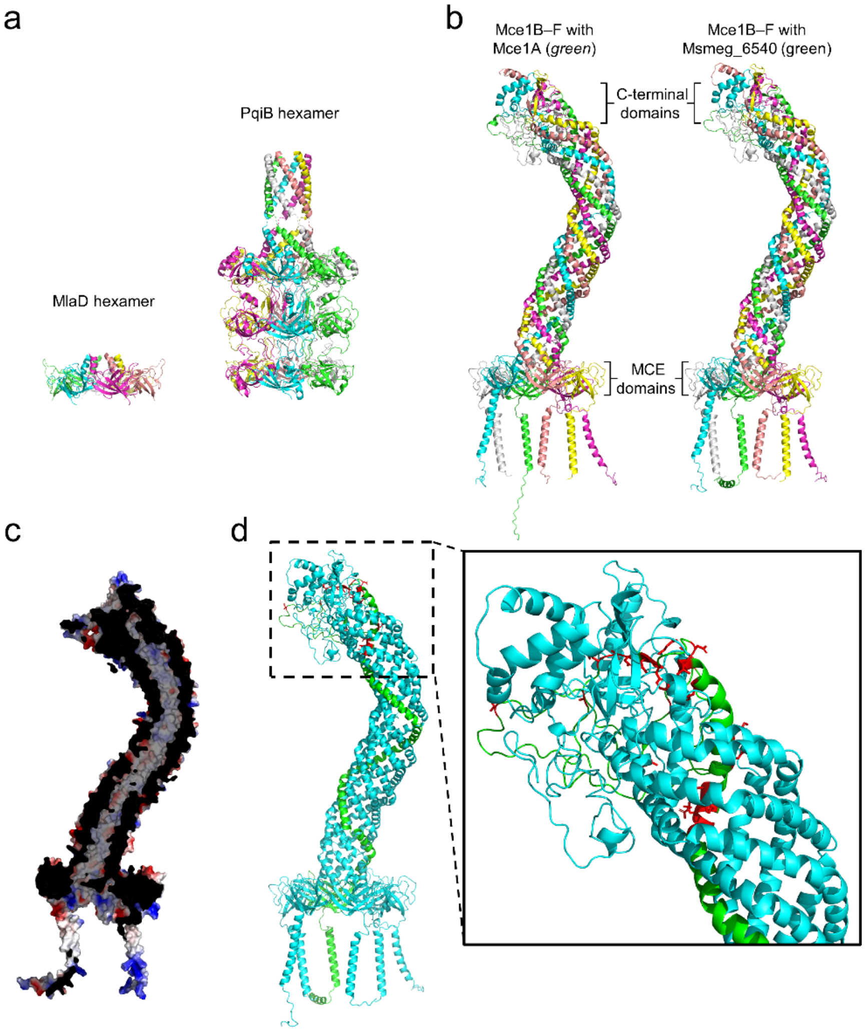


**Supplementary Figure 7. Mce1B–F form putative heterohexamers with Mce1A or MSMEG_6540. a** Structures of the hexamer of MlaD periplasmic domains (left, single MCE domain per protomer, PDB: 5UW2) and the hexamer of PqiB periplasmic domains (right, three MCE domains per protomer, PDB: 5UVN)[2]. **b** AlphaFold2 structural models of putative heterohexamers formed by either Mce1A or MSMEG_6540 with Mce1B–F. Long unstructured C-terminal tails of Mce1C, Mce1D and Mce1F in the models were removed for the purpose of clarity. *green*: Mce1A or MSMEG_6540, *yellow*: Mce1B, *salmon*: Mce1C, *cyan*: Mce1D, *magenta*: Mce1E, *grey*: Mce1F. **c** Cross-section view of the electrostatic surface map of the hexamer formed by MSMEG_6540 and Mce1B–F, revealing a hydrophobic tunnel spanning the entire assembly. **d** The AlphaFold2 structural model of the hexamer formed by MSMEG_6540 (*green*) and Mce1B–F (cyan) with residues in the C-terminal domain of MSMEG_6540 differing from those on Mce1A highlighted in *red*.

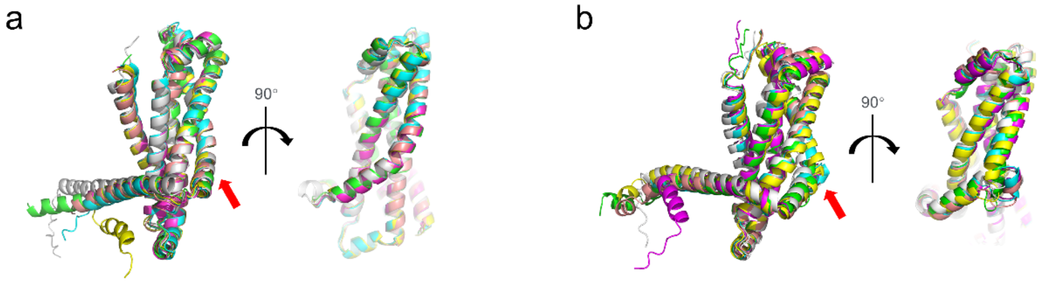


c

```
              1                                                   50
YrbE7B        ~~MTA....S PYVPTLLAPW VRFGQRASKP IVRLGHMLVF FVRAVAAIPI
YrbE1B        ~~MSTVQVLR SRFPRAFSRS SEIAATPARF LDSMGHVAWF VVQAIVHVPH
YrbE3B        ~~~~~~~~~~ ~~~~~~~~~M SNRLDRVRAA WDGIGVQTQF YARAIGSIGD
YrbE4B        MRKKEKPMSY DATL....RF RRLFRGVPKT VDTVGEQALF YGETMRYLPN
YrbE5B        ~~~~~~MVVR TLHP....RL TRELRRPVDV AGRLGDHTSF YGRALAGTPH
YrbE5bisB     ~~~~~MTILR STYP....RL TRQFNKPVAT LSRVGDHTLF YLRAIAGTPH

              51                                                  100
YrbE7B        VLRHYRGEFV RLLSDIAWGN GSLVVGGGTA GVAVVLGITV GALVGIEGYN
YrbE1B        AFRHYRRESL RLVAEIGMGT GAMAVIGGTV AIIGFVTLSA GSLIAIQGFA
YrbE3B        AAMNYRTELI RLIATMGLGA GALAVVGGTV AIVGFLTMTT GALVAVQGYN
YrbE4B        AFTRYRKETI RLVAEMTMGA GALVMIGGTV GVAAFLTLAS GGVIAVQGYS
YrbE5B        AAMHYRREVI RLIAEISMGA GTLAMIGGTV VIVGFLTLAA GGTLAVQGYS
YrbE5bisB     AALHYRKELV RLIAEISMGA GTLAMIGGTV VIVGFLTLAT GGVLAIQGYS

              101                                                 150
YrbE7B        FLDLLGLGPA TGIISSLVNT RELAPIAASL AFATQAGCRF TAQLGSMRIA
YrbE1B        SLGNIGVEAF TGFFAALANI RVVAPVVTGQ ALAATVGAGA TAELGAMRIS
YrbE3B        QFASVGVEAL TGFASAFFNV RLIAPGTTAV ALSATIGAGA TAQLGAMRIN
YrbE4B        SLGNIGIEAL TGFLSAFLNV RIVAPVIAGI ALAATIGAGA TAQLGAMRVA
YrbE5B        SLGNIGIEAL TGFLAAFINV RISAPVVAGI GLAATFGAGV TAQLGAMRIN
YrbE5bisB     SLGNIGIEAL TGFLSAFINV RIAAPVVAGI GLAATFGAGV TAQLGAMRIN

              151                                                 200
YrbE7B        EEIDALDSIA IRPIPYLVTT RLMAAVVAVI PLYVMCLAVS YLTTQTVVFL
YrbE1B        EEVDALEVMG IKSISYLVST RIMAGAIVII PLYAMAILLS FMSAQLVTTI
YrbE3B        EEIDALEVIG IRSVSYLAST RVAAGVIVVI PLYCVAVIMS FISSRLGTTV
YrbE4B        EEIDAVESMA VHAVSYLVST RLLAGLIAIV PLYSLSVLAA FFAARFTTVF
YrbE5B        EEIDALESMG IRPVEYLVST RIVAGMVAIT PLYAVAVILS FAASNLTTVV
YrbE5bisB     EEIDALESMA IRPVEYLVST RIVAGMIAIT PLYAIAVILS FLASQFTTVV

              201                                                 250
YrbE7B        ISGGSTGSYL HYFTLMLSGQ DILYSLLKTI IFVWIASTVQ CYYGFYASGG
YrbE1B        FYSQSVGTYE HYFHTFLRVD DVMWSFLEVI IMSVIVMLNH CYFGYFASGG
YrbE3B        IYGQGSGVYD HYFNTFLNTN DLLWSFGQSV AITVVIMLVH TYYGYTASGG
YrbE4B        INGQSAGLYD HYFNTFLIPS DLLWSFLQAI VMAIAVMLVH TYYGYNASGG
YrbE5B        MFGQSAGLYN HYFSTFLNPV DLLWSFLQAI LMAIAILLVH TYFGFFAAGG
YrbE5bisB     LLGQSGGLYN HYFDTFLNPI DLLWSFLQAI LMAITILLIH TYFGYFASGG

              251                                     291
YrbE7B        PVGVGVAAGH AMRASITVVI IVNMLLTMAL WSVDSGARFG G
YrbE1B        AVGVGEAVGR SMRTSLIAIV LVVLLASLAL YGTDPNFNLT V
YrbE3B        PAGVGEAVGR AVRTSLIVAA IVVVMISLAL YGQSGNFHLA G
YrbE4B        PVGVGIAVGQ AVRTSLIVVV TITLFISLAV YGASGNFNLS G
YrbE5B        PSGVGVAVGN AVRTSLIVVI AVTLLVSLAV YGSNGNFNLS G
YrbE5bisB     PSGVGVAVGN AVRTSLIVVV SVTLLVSLAV YGSNGNFNLS G
```

**Supplementary Figure 8. The putative MSMEG_0959 binding motif may be conserved across YrbEBs.** **a** Overlay of Alphafold2 structural models of six YrbEAs in *M. smegmatis*. **b** Overlay of Alphafold2 structural models of six YrbEBs in *M. smegmatis*. In **a** and **b**, the region where YrbE1B is predicted to interact with MSMEG_0959, and the corresponding region in YrbEAs, are indicated with red arrows. **c** Sequence alignment of six YrbEBs in *M. smegmatis*. The putative helix that is required for the interaction of YrbE1B with MSMEG_0959 is boxed in *red*.

**Supplementary Table 1**. Percentage sequence identity of Mce1A and its homologs with other Mce proteins in *M. smegmatis*.

| % seq identity | Mce1A | 6540 | 5818 | Mce1B | Mce1C | Mce1D | Mce1E | Mce1F | Mce3A | Mce4A | Mce5A | Mce5b isA | Mce7A |
|---|---|---|---|---|---|---|---|---|---|---|---|---|---|
| Mce1A | | 79.8 | 59.8 | 14.9 | 16.3 | 18.4 | 20.0 | 14.7 | 28.1 | 28.7 | 31.9 | 29.6 | 16.3 |
| Msmeg_6540 | | | 57.7 | 16.7 | 17.1 | 14.5 | 19.6 | 18.7 | 26.1 | 26.8 | 26.9 | 28.2 | 17.9 |
| Msmeg_5818 | | | | 17.8 | 16.4 | 13.5 | 22.0 | 19.1 | 28.2 | 28.5 | 30.8 | 30.3 | 18.9 |

**Supplementary Table 2**. Area of interfaces between each pair of TMD and NBD protomer within different ABC transporters with heterodimeric TMDs. PDB ID: $LptB_2FG$[3], 6MHZ; $LolCD_2E$[4], 7ARM; $SugABC_2$[5], 7CAG; $MalFGK_2$[6], 2R6G; $AlgM1M2S_2$[7], 4TQU. For YrbE1A/B-MceG, the AlphaFold2 model[8,9] was used for the interface analysis.

| **ABC transporter** | **YrbE1A/B-MceG** | | **$LptB_2FG$** | | **$LolCD_2E$** | | **$SugABC_2$** | | **$MalFGK_2$** | | **$AlgM1M2S_2$** | |
|---|---|---|---|---|---|---|---|---|---|---|---|---|
| TMD-NBD | A-G | B-G | F-B | G-B | C-D | E-D | A-C | B-C | F-K | G-K | M1-S | M2-S |
| Interface area* (Å$^2$) | 699 | 837 | 807 | 1081 | 1286 | 1022 | 756 | 1053 | 775 | 1024 | 836 | 797 |

* Calculated using the interface analysis function in ChimeraX 1.4.

**Supplementary Table 3**. *M. smegmatis* strains used in this study.

| **Strains used** | **Source** |
|---|---|
| mc$^2$155 | lab collection |
| mc$^2$155 Δ*mce1R* | this study |
| mc$^2$155 Δ*mceG* | this study |
| mc$^2$155 Δ*mce1A* | this study |
| mc$^2$155 Δ*MSMEG_6540* | this study |
| mc$^2$155 Δ*mce1B* | this study |
| mc$^2$155 Δ*mce1C* | this study |
| mc$^2$155 Δ*mce1D* | this study |
| mc$^2$155 Δ*mce1E* | this study |
| mc$^2$155 Δ*mce1F* | this study |
| mc$^2$155 Δ*yrbE1A/B* | this study |
| mc$^2$155 Δ*mce1A*Δ*MSMEG_6540* | this study |

**Supplementary Table 4**. Plasmids used in this study for protein expression and gene deletion in *M. smegmatis*.

| plasmid | description | source |
|---|---|---|
| pJEB402 | an integrative vector containing an *attP* site; express genes under an MOP promoter; $Kan^R$ | Lee & Hatfull[10] |
| pJEB402*his₆-yrbE1A* | to express YrbE1A with an N-terminal His-tag under an MOP promoter; $Kan^R$ | this study |
| pJEB402*yrbE1A/B* | to express YrbE1A/B under an MOP promoter; $Kan^R$ | this study |
| pJEB402*mceG* | to express MceG under an MOP promoter; $Kan^R$ | this study |
| pJEB402*mce1A* | to express Mce1A under an MOP promoter; $Kan^R$ | this study |
| pJEB402*MSMEG_6540* | to express MSMEG_6540 under an MOP promoter; $Kan^R$ | this study |
| pJEB402*mce1B* | to express Mce1B under an MOP promoter; $Kan^R$ | this study |
| pJEB402*mce1C–F-mam1A–D* | to express Mce1C–F and Mam1A–D under an MOP promoter; $Kan^R$ | this study |
| pJEB402*mce1B–F-mam1A–D* | to express Mce1B–F and Mam1A–D under an MOP promoter; $Kan^R$ | this study |
| pJEB402*mce1C* | to express Mce1C under an MOP promoter; $Kan^R$ | this study |
| pJEB402*mce1D* | to express Mce1D under an MOP promoter; $Kan^R$ | this study |
| pJEB402*mce1E* | to express Mce1E under an MOP promoter; $Kan^R$ | this study |
| pJEB402*mce1E–F-mam1A–D* | to express Mce1E–F and Mam1A–D under an MOP promoter; $Kan^R$ | this study |
| pJEB402*mce1F* | to express Mce1F under an MOP promoter; $Kan^R$ | this study |
| pJEB402*mam1A–D* | to express Mam1A–D under an MOP promoter; $Kan^R$ | this study |
| pJEB402*mce1F-mam1A–D* | to express Mce1F and Mam1A–D under an MOP promoter; $Kan^R$ | this study |
| pMV306hsp | an integrative vector containing an *attP* site; express genes under an *hsp60* promoter; $Kan^R$ | Andrew et al.[11] |
| pMV306hsp*his₆-yrbE1A* | to express YrbE1A with an N-terminal His-tag under an *hsp60* promoter; $Kan^R$ | this study |
| pMV306hsp*his₆-yrbE1B* | to express YrbE1B with an N-terminal His-tag under an *hsp60* promoter; $Kan^R$ | this study |
| pMV306hsp$MSMEG_0959_{AGG3P}$ | to express $MSMEG\_0959_{AGG3P}$ under an *hsp60* promoter; $Kan^R$ | this study |

| | | |
|---|---|---|
| pMV306hsp*MSMEG_0959*$_{AGG3P}$-*FLAG+his*$_6$-*yrbE1B* | to express MSMEG_0959$_{AGG3P}$ with a C-terminal FLAG-tag and YrbE1B with an N-terminal His-tag under an *hsp60* promoter; Kan$^R$ | this study |
| pMV306hsp*Rv0513* | to express Rv0513 under an *hsp60* promoter; Kan$^R$ | this study |
| pMV306hsp*Rv0513-FLAG +his*$_6$-*yrbE1B* | to express Rv0513 with a C-terminal FLAG-tag and YrbE1B with an N-terminal His-tag under an *hsp60* promoter; Kan$^R$ | this study |
| pMV306hsp*MSMEG_0959* | to express MSMEG_0959 under an *hsp60* promoter; Kan$^R$ | this study |
| pMV306hsp*MSMEG_0959-FLAG+his*$_6$-*yrbE1B* | to express MSMEG_0959 with a C-terminal FLAG-tag and YrbE1B with an N-terminal His-tag under an *hsp60* promoter; Kan$^R$ | this study |
| pGOAL17 | a vector used as the template for the *lacZ-sacB* cassette under a $P_{Ag85}$ promoter for screening and negative selection; Amp$^R$ | Parish & Stoker[12] |
| pYUB854 | a vector lacking mycobacterial replication origin; Hyg$^R$ | Bardarov et al.[13] |
| pYUB854*53UTR*$_{mce1R}$+*lacZ-sacB* | to achieve unmarked deletion of *mce1R*; Hyg$^R$ | this study |
| pYUB854*53UTR*$_{mceG}$+*lacZ-sacB* | to achieve unmarked deletion of *mceG*; Hyg$^R$ | this study |
| pYUB854*53UTR*$_{mce1A}$+*lacZ-sacB* | to achieve unmarked deletion of *mce1A*; Hyg$^R$ | this study |
| pYUB854*53UTR*$_{MSMEG\_6540}$+*lacZ-sacB* | to achieve unmarked deletion of *MSMEG_6540*; Hyg$^R$ | this study |
| pYUB854*53UTR*$_{mce1B}$+*lacZ-sacB* | to achieve unmarked deletion of *mce1B*; Hyg$^R$ | this study |
| pYUB854*53UTR*$_{mce1C}$+*lacZ-sacB* | to achieve unmarked deletion of *mce1C*; Hyg$^R$ | this study |
| pYUB854*53UTR*$_{mce1D}$+*lacZ-sacB* | to achieve unmarked deletion of *mce1D*; Hyg$^R$ | this study |
| pYUB854*53UTR*$_{mce1E}$+*lacZ-sacB* | to achieve unmarked deletion of *mce1E*; Hyg$^R$ | this study |
| pYUB854*53UTR*$_{mce1F}$+*lacZ-sacB* | to achieve unmarked deletion of *mce1F*; Hyg$^R$ | this study |
| pYUB854*53UTR*$_{yrbE1A/B}$+*lacZ-sacB* | to achieve unmarked deletion of *yrbE1A/B*; Hyg$^R$ | this study |

**Supplementary Table 5**. Plasmids used in this study for protein expression in *E. coli*.

| plasmid | description | source |
|---|---|---|
| pET28b(+) | a high-copy vector; to express genes under an IPTG-inducible T7 promoter; pBR322 ori; $Kan^R$ | Novagen |
| pET22/42 | a high-copy vector; made of the backbone of pET22b(+) and the multiple cloning site of pET42a(+); to express genes under an IPTG-inducible T7 promoter; pBR322 ori; $Amp^R$ | lab collection |
| pCDFDuet1 | a high-copy vector; contains two multiple cloning sites to express genes under two separate IPTG-inducible T7 promoters; CloDF13 ori; $Spec^R$ | Novagen |
| pET28b*his*$_6$*-yrbE1A* | to express YrbE1A with an N-terminal His-tag under an IPTG-inducible T7 promoter; pBR322 ori; $Kan^R$ | this study |
| pET28b*his*$_6$*-yrbE1B* | to express YrbE1B with an N-terminal His-tag under an IPTG-inducible T7 promoter; pBR322 ori; $Kan^R$ | this study |
| pET22/42*yrbE1A*-(*his*$_6$-*yrbE1B*) | to express YrbE1A and YrbE1B with an N-terminal His-tag using gene sequences optimized for *E. coli* codon usage under an IPTG-inducible T7 promoter; pBR322 ori; $Amp^R$ | this study |
| pET22/42(*his*$_6$-*yrbE1A*)-*yrbE1B* | to express YrbE1A with an N-terminal His-tag and YrbE1B using gene sequences optimized for *E. coli* codon usage under an IPTG-inducible T7 promoter; pBR322 ori; $Amp^R$ | this study |
| pET22/42*yrbE1A*-(*his*$_6$-*yrbE1B*$_{GS}$) | to express YrbE1A and YrbE1B (the helix mutant) with an N-terminal His-tag using gene sequences optimized for *E. coli* codon usage under an IPTG-inducible T7 promoter; pBR322 ori; $Amp^R$ | this study |
| pCDFDuet1*mceG* | to express MceG under an IPTG-inducible T7 promoter; CloDF13 ori; $Spec^R$ | this study |
| pCDFDuet1*mceG*$_{K43A}$ | to express $MceG_{K43A}$ under an IPTG-inducible T7 promoter; CloDF13 ori; $Spec^R$ | this study |
| pCDFDuet1*mceG-FLAG* | to express MceG with a C-terminal FLAG-tag under an IPTG-inducible T7 promoter; CloDF13 ori; $Spec^R$ | this study |
| pCDFDuet1*MSMEG_0959-FLAG* | to express MSMEG_0959 with a C-terminal FLAG-tag under an IPTG-inducible T7 promoter; CloDF13 ori; $Spec^R$ | this study |
| pCDFDuet1*mceG-FLAG+MSMEG_0959* | to express MceG with a C-terminal FLAG-tag and MSMEG_0959 under two separate IPTG-inducible T7 promoters; CloDF13 ori; $Spec^R$ | this study |
| pCDFDuet1*mceG-FLAG+MSMEG_0959-FLAG* | to express MceG with a C-terminal FLAG-tag and MSMEG_0959 with a C-terminal FLAG-tag under two separate IPTG-inducible T7 promoters; CloDF13 ori; $Spec^R$ | this study |

| | | |
|---|---|---|
| pCDFDuet1*mceG-FLAG+MSMEG_0959*$_{AGG3P}$-*FLAG* | to express MceG with a C-terminal FLAG-tag and MSMEG_0959$_{AGG3P}$ with a C-terminal FLAG-tag under two separate IPTG-inducible T7 promoters; CloDF13 ori; $Spec^R$ | this study |

**Supplementary Table 6**. Primers used in this study.

| name | sequence (5' to 3') | function |
|---|---|---|
| 1R5UTRF | AGCTACTAGTGACCTGTTCGGACTCAAGCACATC | to construct the suicide plasmid for *mce1R* deletion by restriction cloning |
| 1R5UTRR | ATATAAGCTTGTGGTCTCCGGTGGTATCTCGGGTG | |
| 1R3UTRF | ACGTAAGCTTCAGATCAGCTCTGGGTGAGC | |
| 1R3UTRR | ATATCCATGGTGCACCGATGCGTTCGAGGACGGCT | |
| G5UTRF | ATATACTAGTTCGGAAGCCGGCATTGAAGGCCTCGA | to construct the suicide plasmid for *mceG* deletion by restriction cloning |
| G5UTRR | AGCTAAGCTTCAAAGATCCTTCCCGCTACGCCTACCA | |
| G3UTRF | AATTAAGCTTGTTTGCCGCGATCAGGCCGGGCCGTCA | |
| G3UTRR | ATATCCATGGATCCGAGAAGGACAGTGACATCGAG | |
| 1A5UTRF | TGGTAATACGACTCA GAAGGCGACGCTGTTCGGAA | to construct the suicide plasmid for *mce1A* deletion by Gibson assembly |
| 1A5UTRR | TTCTCCC CGCTACACCGTCAGGTTGAAG | |
| 1A3UTRF | GTGTAGCG GGGAGAACACGATCAACCCAT | |
| 1A3UTRF | TATTTCGGACGGTTG CCGTTGAGCTTCAAGGTCA | |
| 65405UTRF | CGTTACTAGTCAAGGCGATGATGGTCTTCTCC | to construct the suicide plasmid for *MSMEG_6540* deletion by restriction cloning |
| 65405UTRR | ATATAAGCTTCGCCGCGGATGAGGCCACCT | |
| 65403UTRF | AGCTAAGCTTCAACCCGTGACTGTCTTGGCT | |
| 65403UTRR | AATTCCATGGCTGCTCGACATCACCGGCA | |
| 1B5UTRF | TGGTAATACGACTCA CCGCCTGGCGTCGATCGACGTCG | to construct the suicide plasmid for *mce1B* deletion by Gibson assembly |
| 1B5UTRR | TGTCCTC ATGGGTTGATCGTGTTCTCCCCCA | |
| 1B3UTRF | CAACCCAT GAGGACACTGCAGGGTTCCGAC | |
| 1B3UTRF | TATTTCGGACGGTTG TCGACGAAGGGCTGCAGGATG | |
| 1C5UTRF | TGGTAATACGACTCA TCGGCATCTTCTCGCTGGTGCT | to construct the suicide plasmid for *mce1C* deletion by Gibson assembly |
| 1C5UTRR | TCCCTGC CTATTTCGGCGTGCACCTACC | |
| 1C3UTRF | CGAAATAG GCAGGGAGGCGTCGAGACATGT | |
| 1C3UTRF | TATTTCGGACGGTTG CCTGGTTGGGCGAGTTGATGTT | |
| 1D5UTRF | TGGTAATACGACTCA AACAAGGTCGCGACGGTGCTCG | to construct the suicide plasmid for *mce1D* deletion by Gibson assembly |
| 1D5UTRR | AGCCTCAT GTCTCGACGCCTCCCTGCCTAT | |
| 1D3UTRF | TCGAGAC ATGAGGCTGCTGAAGGGTTTCC | |
| 1D3UTRF | TATTTCGGACGGTTG AGGTCGAGCTTGAGCGACACGT | |
| 1E5UTRF | TGGTAATACGACTCA GTTTGGCATTGTTCGTCAACGC | to construct the suicide plasmid for *mce1E* deletion by Gibson assembly |
| 1E5UTRR | CCCTTTC TCAGCCTGCTCCTGCTTCAGCGGG | |
| 1E3UTRF | CAGGCTGA GAAAGGGGGGAGTGAGATGCTG | |
| 1E3UTRF | TATTTCGGACGGTTG CACTGCGAGGCGGGCAGGAAGC | |
| 1F5UTRF | ATATACTAGTAGCATGACGCTGTACGTGCAGA | to construct the suicide plasmid for *mce1F* deletion by restriction cloning |
| 1F5UTRR | ATATAAGCTTCTCACTCCCCCCTTTCGAC | |
| 1F3UTRF | ATATAAGCTTAGGAGATGACGGATGGAAGG | |
| 1F3UTRF | ATATCCATGGTCCGGTTACGCGCGTCCCA | |
| y15UTRF | AATTACTAGTGAAACTGCGTGCGCTGTCGT | to construct the suicide plasmid for *yrbE1A/B* deletion by restriction cloning |
| y15UTRR | ATATAAGCTTAGGTGCCCTTCCTGGACGTGATC | |
| y13UTRF | ATATAAGCTTCTGACGGTGTAGCGCCATGAC | |
| y13UTRR | AATTCCATGGGTACACATACGGGTTGCC | |
| lZsBF | CGCGGCCGCAATTAACCCTCACTAAAGGATC<br>TTAATTAAAGCCCGCTCATTAGGCGGG | to clone the *lacZ-sacB* cassette using pGOAL17 as the template for insertion into suicide plasmids by Gibson assembly |
| lZsBR | GGCCGCATAATACGACTCACTATAGGGATC<br>TTAATTAAGCGGCCGCGGTACCAA | |
| y1AF | ATCACATATGACGGCGTCGACCGATG | to construct pET28b*his*$_6$*-yrbE1A* by restriction cloning |
| y1AR | ATGTAAGCTTTCAGCGCCCTGTCCCGAAC | |
| y1BF | ATCACATATGAGTACTGTTCAGGTTCTCCGCT | to construct pET28b*his*$_6$*-yrbE1B* by restriction cloning |
| y1BR | ATGTAAGCTTCTACACCGTCAGGTTGAAGTTCG | |
| opt1NBF | AATTCATATGACTGCGTCTACAGAC | to construct pET22/42*yrbE1A*-(*his*$_6$*-yrbE1B*) by restriction |
| opt1NBR | AGCTAAGCTTCTATACAGTAAGATTAAAGTTAGG | |

| | | |
|---|---|---|
| | | cloning with *yrbE1A*-(*his*$_6$-*yrbE1B*) cloned from a synthesized *mce1* operon optimized for *E. coli* codon usage |
| opt1BF | ACCATGAGTACAGTACAGGTGTTGC | to construct pET22/42*yrbE1A*/*B* by site-directed mutagenesis (SDM) using pET22/42*yrbE1A*-(*his*$_6$-*yrbE1B*) as the template |
| opt1BR | TGTACTCATGGTATATCTCCTTCTTAAATC | |
| opt1NAF | CACCATCATCACCACAGCAGCGGCACTGCGTCTACAGACGGGTT | to construct pET22/42(*his*$_6$-*yrbE1A*)-*yrbE1B* by SDM using pET22/42*yrbE1A*/*B* as the template |
| opt1NAR | GTGATGATGGTGATGGCTGCTGCCCATATGTATATCTCCTTCTTAAAG | |
| opt1NBGSF | GTGGTAGCAGCGGTGGGGCGATGGCTGTCATC | to construct pET22/42*yrbE1A*-(*his*$_6$-*yrbE1B*$_{GS}$) using pET22/42*yrbE1A*-(*his*$_6$-*yrbE1B*) as the template by SDM |
| opt1NBGSR | CGCTGCTACCACCCAAAGATTCACGACGGTAGT | |
| CDFGF | TATACCATGGCGTCCAAATCGACGT | to construct pCDFDuet1*mceG* by restriction cloning |
| CDFGR | ATATAAGCTTCACGCCTGCTTGGGCA | |
| GKAF | ACCGGCGCGTCCGTGTTCCTGAAGTCGCTG | to construct pCDFDuet1*mceG*$_{K43A}$ using pCDFDuet1*mceG* as the template by SDM |
| GKAR | GGACGCGCCGGTACCGGAGGGGCCCAGCA | |
| GflF | GACTACAAGGACGATGACGACAAGTGAAGCTTGCGGCCGCATAA | to add a FLAG-tag sequence to *mceG* at the first multiple cloning site of pCDFDuet1 by SDM |
| GflR | GTCCTTGTAGTCACCGCTGCTACCCGCCTGCTTGGGCACCTCGA | |
| CDF959F | AATTCATATGACTGAACCCGCAGGCCACGA | to insert *MSMEG_0959* into the second multiple cloning site of pCDFDuet1 by restriction cloning |
| CDF959R | TATACCTAGGTCACGGCGTGCGCTCTTCCACC | |
| AGPF | CGGAGCTCACCCCGGTCCCGCGCGGCCGGACC | to convert WT *MSMEG_0959* to *MSMEG_0959*$_{AGG3P}$ on different plasmids by SDM |
| AGPR | GGGTGAGCTCCGGGAGCGGACGCGCCGACTGCCACTT | |
| Jy1F | CGGATCCAGCTGCAG GAAGGGCACCTGTGACGG | to construct pJEB402*yrbE1A/B* by Gibson assembly |
| Jy1R | GTGCGGCCGCGGTAC CTACACCGTCAGGTTGAAGT | |
| JGF | CGGATCCAGCTGCAG TAAGGAGATATACCATGGGC | to construct pJEB402*mceG* with *mceG* cloned from pCDFDuet1*mceG* as the template by Gibson assembly |
| JGF | GTGCGGCCGCGGTAC TCACGCCTGCTTGGGCACCT | |
| J1AF | CGGATCCAGCTGCAG AGGAAGGAGCGCCATGACCGAGCCTCCAGCGC | to construct pJEB402*mce1A* using Gibson assembly |
| J1AR | ACGTCGACATCGATA CTCATGGGTTGATCGTGTTCT | |
| J6540F | ATATGAATTCAGGAGAAGCGCCGTGGCCTCATCCGCGGCGT | to construct pJEB402*MSMEG_6540* by restriction cloning |
| J6540R | ATATAAGCTTTCACGGGTTGATGGTGTTCT | |
| Jm1F | CGGATCCAGCTGCAG AGGAGATGACGGATGGAAGGA | to construct pJEB402*mam1A–D* by Gibson assembly |
| Jm1R | ACGTCGACATCGATACTACAGCTGCGGCGAGAGGTC | |
| J1BF | CGGATCCAGCTGCAG AGGAAGGAGACCCATGAGTATCAAAGGCACGCTTT | to construct pJEB402*mce1B* by Gibson assembly; J1BF was also used together with Jm1R to construct pJEB402*mce1B–F-mam1A–D* |
| J1BR | ACGTCGACATCGATA TCATTTCGGCGTGCACCTACC | |
| J1CF | CGGATCCAGCTGCAG AGGAAGGAGCGAAATGAGGACACTGCAGGGTTCC | to construct pJEB402*mce1C* by Gibson assembly; J1CF was also used together with Jm1R to construct pJEB402*mce1C–F-mam1A–D* |
| J1CR | ACGTCGACATCGATA CTATCTCGACTGCGAACCCGG | |

| | | |
|---|---|---|
| J1DF | CGGATCCAGCTGCAG AGGAAGGAGAGACATGTCAACGATTTTCAACATCC | to construct pJEB402*mce1D* by Gibson assembly; J1DF was also used together with Jm1R to construct pJEB402*mce1D–F-mam1A–D* |
| J1DR | ACGTCGACATCGATA TCAGCCTGCTCCTGCTTCAGC | |
| J1EF | CGGATCCAGCTGCAG AGGAAGGAGGCTGATGAGGCTGCTGAAGGGTTTCC | to construct pJEB402*mce1E* by Gibson assembly; J1EF was also used together with Jm1R to construct pJEB402*mce1E–F-mam1A–D* |
| J1ER | ACGTCGACATCGATA TCTCACTCCCCCCTTTCGACCAG | |
| J1FF | CGGATCCAGCTGCAG AAAGGGGGGAGTGAGATGCT | to construct pJEB402*mce1F* by Gibson assembly; J1FF was also used together with Jm1R to construct pJEB402*mce1F-mam1A–D* |
| J1FR | GTGCGGCCGCGGTAC TGTGTTCCTGCTGCTGATCG | |
| pETRBSF | TCGA<u>GAATTC</u>TTAACTTTAAGAAGGAGATATAC | to construct pMV306hsp*his*$_6$-*yrbE1A* and pMV306hsp*his*$_6$-*yrbE1B* by restriction cloning with *his*$_6$-*yrbE1A* and *his*$_6$-*yrbE1B* cloned from pET28b*his*$_6$-*yrbE1A* or pET28b*his*$_6$-*yrbE1B* respectively; used together with y1AR or y1BR |
| M959F | ATAT<u>GAATTC</u>AGGAGAGACCGTGACTGAACC | to construct pMV306hsp*MSMEG_0959* by restriction cloning |
| M959R | AATT<u>AAGCTT</u>CACGGCGTGCGCTCTTCC | |
| 959flR | GTCCTTGTAGTCACCGCTGCTACCCGGCGTGCGCTCTTCCACCA | to add a FLAG-tag sequence to *MSMEG_0959* by SDM; used together with M959flF, M959flN1BF or CDF959flF. |
| M959flF | GACTACAAGGACGATGACGACAAGTGAAGCTTATCGATGTCGACG | to add a FLAG-tag sequence to *MSMEG_0959* on pMV306hsp*MSMEG_0959* by SDM |
| M959flN1BF | GACTACAAGGACGATGACGACAAGTGACTTAACTTTAAGAAGGAGATATACC | to add a FLAG-tag sequence to *MSMEG_0959* on pMV306hsp*MSMEG_0959+his*$_6$-*yrbE1B* by SDM |
| CDF959flF | GACTACAAGGACGATGACGACAAGTGACCTAGGCTGCTGCCACC | to add a FLAG-tag sequence to *MSMEG_0959* at the second multiple cloning site of pCDFDuet1*MSMEG_0959* by SDM |
| MRv0513F | ATAT<u>GAATTC</u>AAGGAGATATACATGACACCAACCGGGGATA | to construct pMV306hsp*Rv0513* by restriction cloning |
| MRv0513R | ATAT<u>AAGCTT</u>CGGCACAAGCCAGGAACAGTT | |
| MRv0513N1BF | GGGATCCAGCTGCAG AAGGAGATATACATGACACCAACCGGGGATA | to construct pMV306hsp*Rv0513 +his*$_6$-*yrbE1B* by Gibson assembly |
| MRv0513N1BR | ATCTCCTTCTTAAAGTTAAG CGGCACAAGCCAGGAACAGTT | |
| Rv0513flF | GACTACAAGGACGATGACGACAAGTGATCGCCCGCTACCGG | to add a FLAG-tag sequence to *Rv0513* on pMV306hsp*Rv0513+his*$_6$-*yrbE1B* by SDM |
| Rv0513flR | GTCCTTGTAGTCACCGCTGCTACCCCGCGCTGGCCCGGCGT | |
| 959L55CF | GATCTGCCAGCGCACCGGCGGTTAC | |

| | | |
|---|---|---|
| 959L55CR | CGCTGGCAGATCGCCATGCCGACACC | to introduce a L55C mutation to MSMEG_0959 by SDM |
| 959G59CF | CACCTGCGGTTACGGCTATGACCTG | to introduce a G59C mutation to MSMEG_0959 by SDM |
| 959G59CR | TAACCGCAGGTGCGCTGCAGGATCGC | |
| 959G60CF | CGGCTGCTACGGCTATGACCTGTGG | to introduce a G60C mutation to MSMEG_0959 by SDM |
| 959G60CR | CCGTAGCAGCCGGTGCGCTGCAGGAT | |
| 959Y61CF | CGGTTGCGGCTATGACCTGTGGATC | to introduce a Y61C mutation to MSMEG_0959 by SDM |
| 959Y61CR | TAGCCGCAACCGCCGGTGCGCTGCAG | |
| 1BE208CF | GTACTGCCACTACTTCCACACGTTC | to introduce a E208C mutation to YrbE1B by SDM |
| 1BE208CR | TAGTGGCAGTACGTGCCGACCGACTG | |
| 1BF211CF | CTACTGCCACACGTTCCTGCGTGTC | to introduce a F211C mutation to YrbE1B by SDM |
| 1BF211CR | GTGTGGCAGTAGTGCTCGTACGTGCC | |
| 1BH212CF | CTTCTGCACGTTCCTGCGTGTCGAC | to introduce a H212C mutation to YrbE1B by SDM |
| 1BH212CR | AACGTGCAGAAGTAGTGCTCGTACGT | |

*Mycobacterium smegmatis* mc$^2$ 155 genomic DNA was used as the template for cloning unless stated otherwise. Restriction sites are underlined. A space is left between the sequence annealed to the template and the sequence used as an overhang for Gibson assembly. SDM, site-directed mutagensis.
